# An explainable deep learning classifier of bovine mastitis based on whole genome sequence data - circumventing the p>>>n problem

**DOI:** 10.1101/2023.03.23.533903

**Authors:** K. Kotlarz, M. Mielczarek, P. Biecek, K. Wojdak-Maksymiec, T. Suchocki, P. Topolski, W. Jagusiak, J. Szyda

## Abstract

The most serious drawback underlying the biological annotation of Whole Genome Sequence data is the p>>n problem, meaning that the number of polymorphic variants (p) is much larger than the number of available phenotypic records (n). Therefore, the major aim of the study was to propose a way to circumvent the problem by combining a LASSO logistic regression model with Deep Learning (DL). That was illustrated by a practical biological problem of classification of cows into mastitis-susceptible or mastitis-resistant, based on genotypes of Single Nucleotide Polymorphisms (SNPs) identified in their WGS. Among several DL architectures proposed via optimisation of DL hyperparameters using the Optuna software, imposed on different SNP sub-sets defined by LASSO logistic regressions with different penalty values, the architecture with 204,642 SNPs was selected as the best one. This architecture was composed of 2 layers with respectively 7 and 46 units per layer as well as respective drop-out rates of 0.210 and 0.358. The classification of the test data set resulted in the AUC=0.750, accuracy=0.650, sensitivity=0.600, and specificity=0.700 was selected as the best model and thus proceeded to genomic and functional annotations. Significant SNPs were selected based on the SHapley Additive exPlanation values transformed to Z-scores to assess the underlying type I-error. These SNPs were annotated to genes. As a final result, a single GO term related to the biological process and thirteen GO terms related to the molecular function were significantly enriched in the gene set that corresponded to the significant SNPs.

**Author Summary:** Our objective is to distinguish between cows that are susceptible and resistant to bovine mastitis by analysing their genomic data. However, we face a significant challenge due to the large number of single nucleotide polymorphisms (SNPs) and limited sample size. To address this challenge, we utilize two methods: feature selection algorithms and deep learning. We experiment with various ways of implementing these techniques and evaluate their performance on a validation set. Our findings reveal that the optimal approach can accurately predict a cow’s susceptibility or resistance status around 65% of the time. Additionally, we employ a technique to identify the most crucial SNPs and their biological functions. Our results indicate that some of these SNPs are related to immune response or protein synthesis pathways, implying that they may affect the cow’s health and productivity.

## Introduction

Due to the development of high-throughput technology, the past few decades have seen a considerable increase in the availability of genomic data [1,2]. Among them, the most common data structure is the whole genome sequence (WGS) that is nowadays available for thousands of individuals representing various species e.g. the European 1+ Million Genomes Initiative for humans [3] or the 1000 Bull Genomes Project for cattle [4]. Effective and efficient computing methods are emerging issues regarding the storage, analysis, and interpretation of this flood of biological information [5,6]. However, the most serious drawback underlying the utilization of WGS data is its statistical nature, the so-called *p*>>*n* problem, meaning that the number of predictors i.e. polymorphic variants (*p*) is much larger than the number of available phenotypic records (*n*) [7]. This impedes the application of standard statistical models, like e.g. regression, unless we decide to split the available predictors into single (oligo) predictor analysis, which is often a case in Genome-Wide Association Studies (GWAS) in which despite the availability of millions of polymorphic variants to test for their association with phenotypes, many single-variant models are applied [8,9], followed by multiple testing correction of the individual hypothesis tests. This is however related to a loss of an important source of information contained in high throughput data, that is the interaction between particular predictors. One possible way around the *p*>>*n* problem is to use models that impose some shrinkage in the estimation of effects – like e.g. mixed models, ridge regression, or LASSO. Another recent trend is to use deep learning (DL) that in many areas offers higher accuracy of classification or prediction. DL has been increasingly used in computational biology, for example, in genomics for identifying regulatory variants [10] or in clinical genetics for predicting an effect of mutations [11] being applied to a whole range of biological material ranging between single cells [12] and tissues [13]. However, despite great flexibility regarding analysed data structures, a critical problem in using DL is the underlying complexity of the algorithms that makes it difficult to interpret the outcome in terms of formally defined statistical hypotheses and consequently to formulate biologically interpretable conclusions.

Therefore, the major aim of the study was to propose a way to circumvent the *p*>>*n* problem by combining a LASSO logistic regression model and DL illustrated by a practical biological problem of classification of cows into mastitis-susceptible or mastitis-susceptible, based on genotypes of Single Nucleotide Polymorphisms (SNPs) identified in their whole genome DNA sequences. This translates to the situation that the number of available SNPs (*p*) vastly exceeds the number of analysed cows (*n*). Furthermore, we tackle the problem of biological explainability of the results on a single SNP level by using SHapley Additive exPlanation values (SHAP) [14].

Bovine mastitis is a disease that is one of the most common disorders in dairy cows [15–17] causing animal welfare problems and economic losses. Mastitis accounts for 38% of all direct costs associated with major production disorders as well as for 70% of the overall losses attributable to mammary tissue injury. That reduces milk production [18] and thus, remains to be the most economically significant disease affecting dairy cattle [19]. The occurrence of bovine mastitis is known to be significantly influenced by several risk factors, including pathogens, host genetics, and environment.

## Results

### Data processing

Due to the poor genome averaged coverage of 3X resulting after the alignment to the reference genome, one cow was removed from the training group. The genome averaged coverage for the remaining 31 individuals in the training group ranged between 7X and 13X, while for the 20 individuals in the test group, it varied between 14 and 37X. After filtering, 16,618,983 SNPs were considered in the downstream analysis. Since SNP calling was performed separately for test and training individuals and genotypes of SNPs that were polymorphic only in one of the data sets, the other group was set to homozygous reference which is the most frequent genotype constellation.

### Optimal DL architecture

The number of SNPs preselected by using the penalised regression approach, varied between 6,665 for the highest penalty expressed by C=0.1 and 1,154,608 for the mildest penalty expressed by C=1.0. Figure 1 presents the numbers of SNPs selected along the decreasing penalty with each subsequent SNP set containing variants from the preceding subsets, (C_n–1_ ⊆ C_n_).

**Fig 1.**
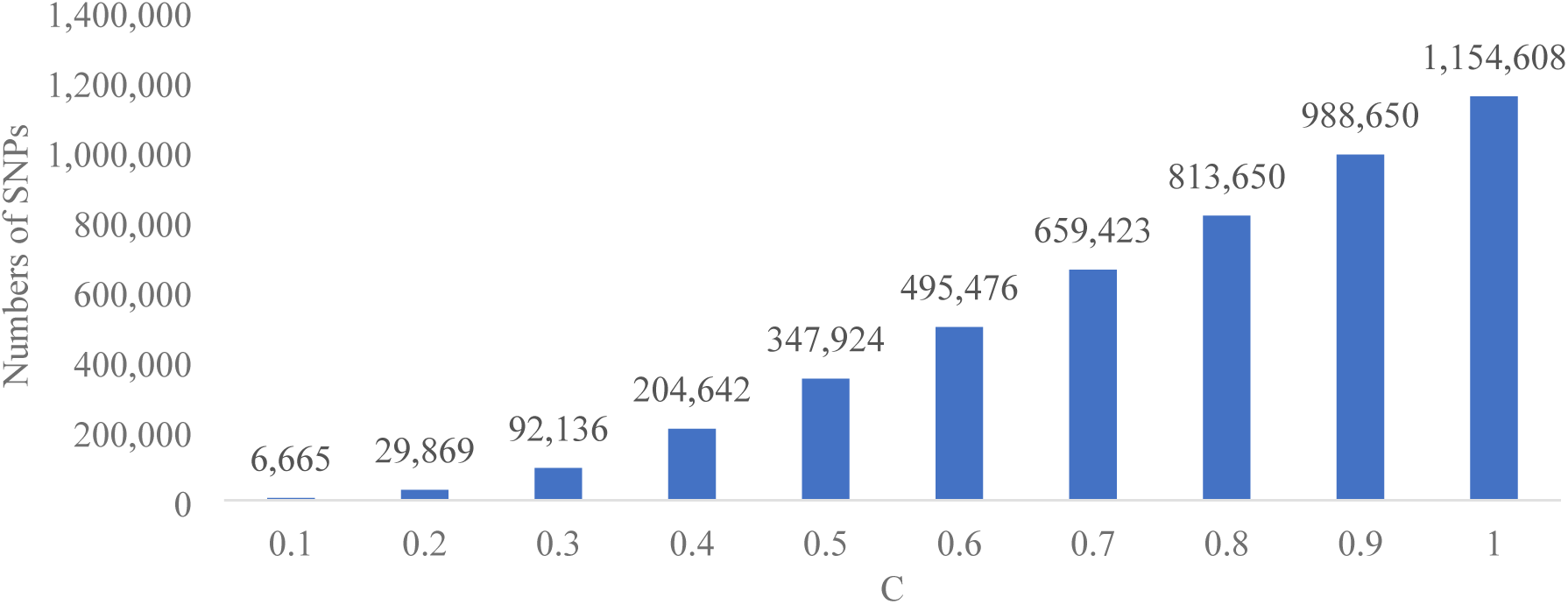
Numbers of SNPs selected using the LASSO logistic regression for each model.

A markedly different DL architecture was selected as the optimal one depending on the SNP-set. The number of layers ranged between one (for C=0.1 and 0.9) and four (C=0.8). For none of the subsets, the maximum allowed number of layers was estimated as the optimal one. The number of units per layer varied between seven and 50, as well as dropout rates were from 0.215 and 0.398 (Table 1).

**Table 1.** The optimal DL architecture estimated for each SNP-set.

| $C = \frac{1}{\lambda}$ | $N$ layers | $N$ units inside each layer | Dropout rate inside each layer | Learning-rate |
| --- | --- | --- | --- | --- |
| 0.1 | 1 | [31] | [0.285] | 3.026e-09 |
| 0.2 | 3 | [32; 36; 37] | [0.302; 0.218; 0.311] | 4.263e-10 |
| 0.3 | 3 | [11; 16; 12] | [0.323; 0.242; 0.243] | 7.147e-09 |
| 0.4 | 2 | [7; 46] | [0.210; 0.358] | 2.328e-11 |
| 0.5 | 2 | [48; 45] | [0.312; 0.250] | 6.700e-12 |
| 0.6 | 3 | [47; 37; 28] | [0.398; 0.278, 0.300] | 6.900e-09 |
| 0.7 | 2 | [10; 18] | [0.215; 0.222] | 7.896e-09 |
| 0.8 | 4 | [35; 35; 26; 13] | [0.323; 0.261, 0.257; 0.327] | 4.268e-10 |
| 0.9 | 1 | [50] | [0.250] | 6.829e-09 |
| 1.0 | 3 | [23; 49; 9] | [0.297; 0.362, 0.365] | 1.698e-09 |

### Classification of individuals

Classification quality expressed by the AUC for each of the estimated DL architectures calculated based on the 4-cross-validation of the training data was summarised in Figure 2A which shows that with AUCs varying between 0.925 (C=0.1) and 1.000 (C=0.9) all the algorithms provided a reasonable classification. Also, the loss for the validation generally decreased with the increasing number of SNPs included in the model, varying between 0.925 for the most parsimonious model with C=0.1 and 0.287 for the model with C=0.9.

**Fig 2.**
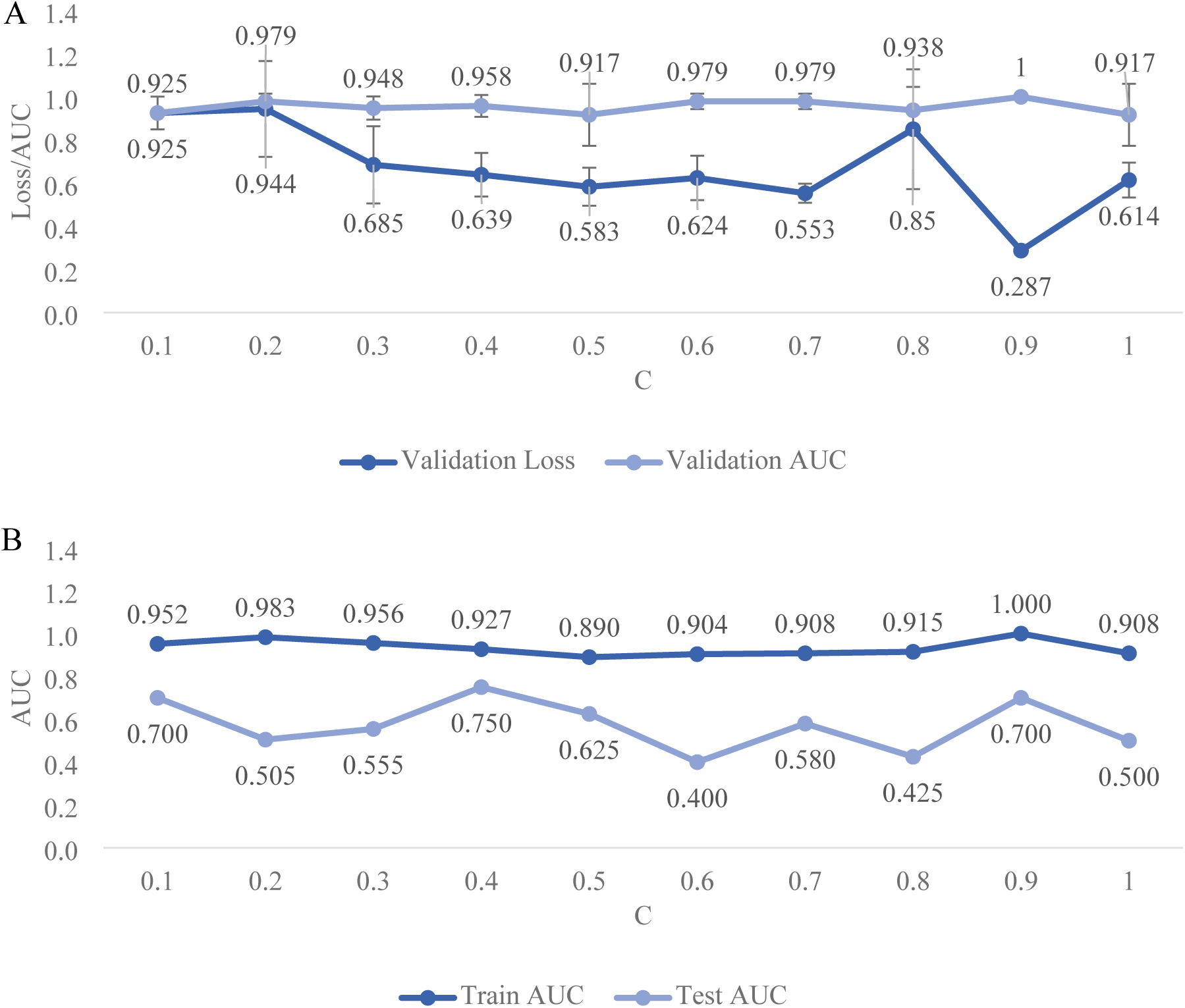
A. AUC and Loss based on the 4-cross-validation of the training data set. B. AUC calculated for the training and test data.

However, when applied to the test data, the classification quality dropped considerably and varied between 0.400 for the SNP set selected under C=0.6 and 0.750 for C=0.4 (Figure 2B). The highest loss in AUC by 0.504 was observed for the SNP set defined by C=0.6, while subsets selected with C=0.1, C=0.4, and C=0.5 were the most robust with 0.252, 0.177 and 0.256 drop in AUC.

### Selection of the best DL architecture

For all the considered SNP-sets, the estimated classification cut-off values differed from the default 0.5 (Figure 3). However, they also considerably differed from each other, from 0.142 (C=0.9) to 0.893 (C=0.2). Cut-off values estimated based on the more complex DL-architectures (0.2, 0.3, and 0.6) were higher than those estimated based on parsimonious architectures. So that the most parsimonious DL-architecture with only one layer, underlying the SNP set obtained under C=0.8 resulted in a very low cut-off value of 0.142. For each DL architecture, the application of the optimal cut-off for the classification of the training data resulted in a higher ACC than using the default cut-off value of 0.5 (Figure 4A). The highest improvement of 0.419 was reached for C=0.4, a SNP-set that based on the default threshold did not even reach the accuracy of a random group assignment (i.e. 0.500). The architecture underlying C=0.9 resulted in a “perfect” accuracy of one, even for the default cut-off, indicating model overfitting. However, when applied to the test data sets the estimated cut-off values did not always result in a better classification. The strongest increase in the accuracy by 0.200 was obtained for SNP-set selected based on C=0.7 (Figure 4B). Moreover, the classification accuracy of test data was much lower as compared to the training data sets and oscillated around 0.500. The highest overall test accuracy of 0.700 was achieved for the data set generated under C=0.9 and the default cut-off. Three SNP-sets (C=0.1, C=0.4, and C=0.7) also resulted in a reasonable accuracy of 0.650 by using estimated cut-off values. The standard errors of the cut-off points did not exceed 0.050, indicating high accuracy of the cut-off values (Figure 3). Figure 4A and Figure 4B, visualise the classification ACC differences obtained for the train and test dataset with the optimal cut-off of the classification algorithms.

**Fig 3.**
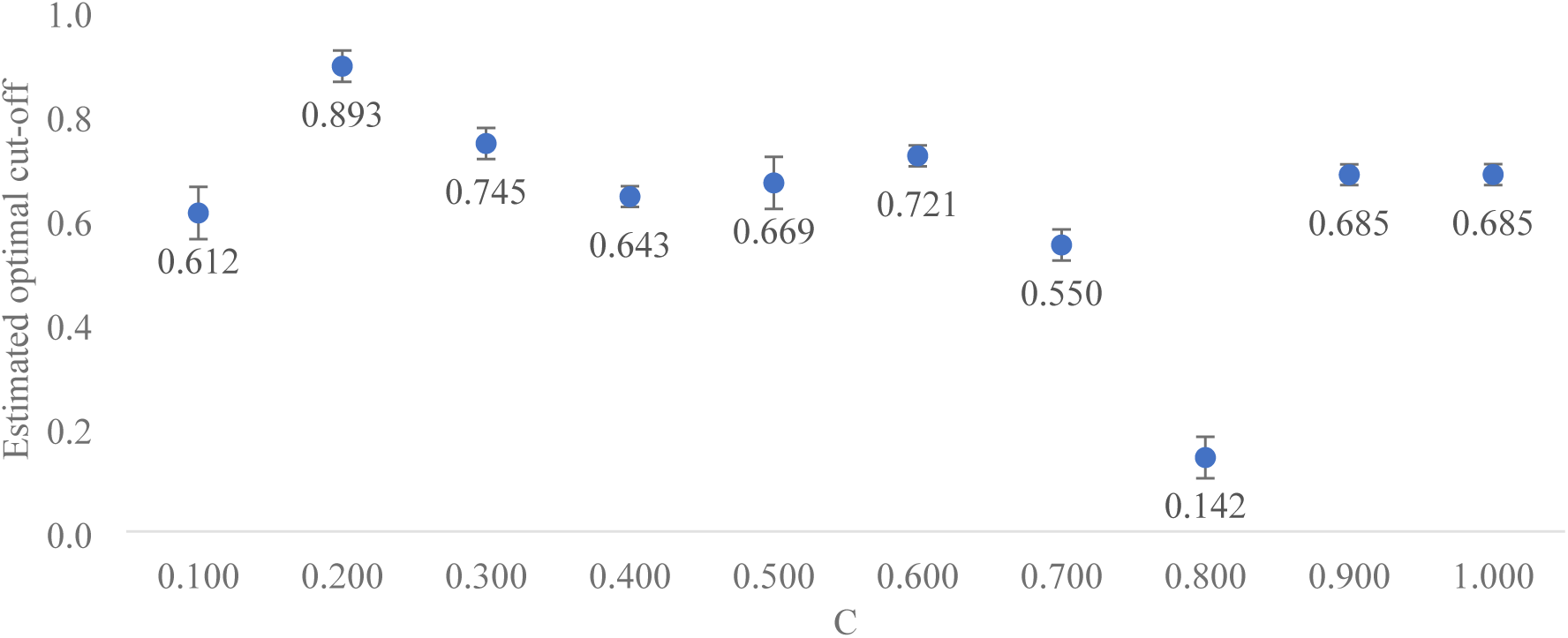
Probability cut-off values for mastitis classification into the susceptible or resistant group estimated based on the optimisation for accuracy metric. The standard deviations for each estimate were calculated using the out-of-the-bag samples.

**Fig 4.**
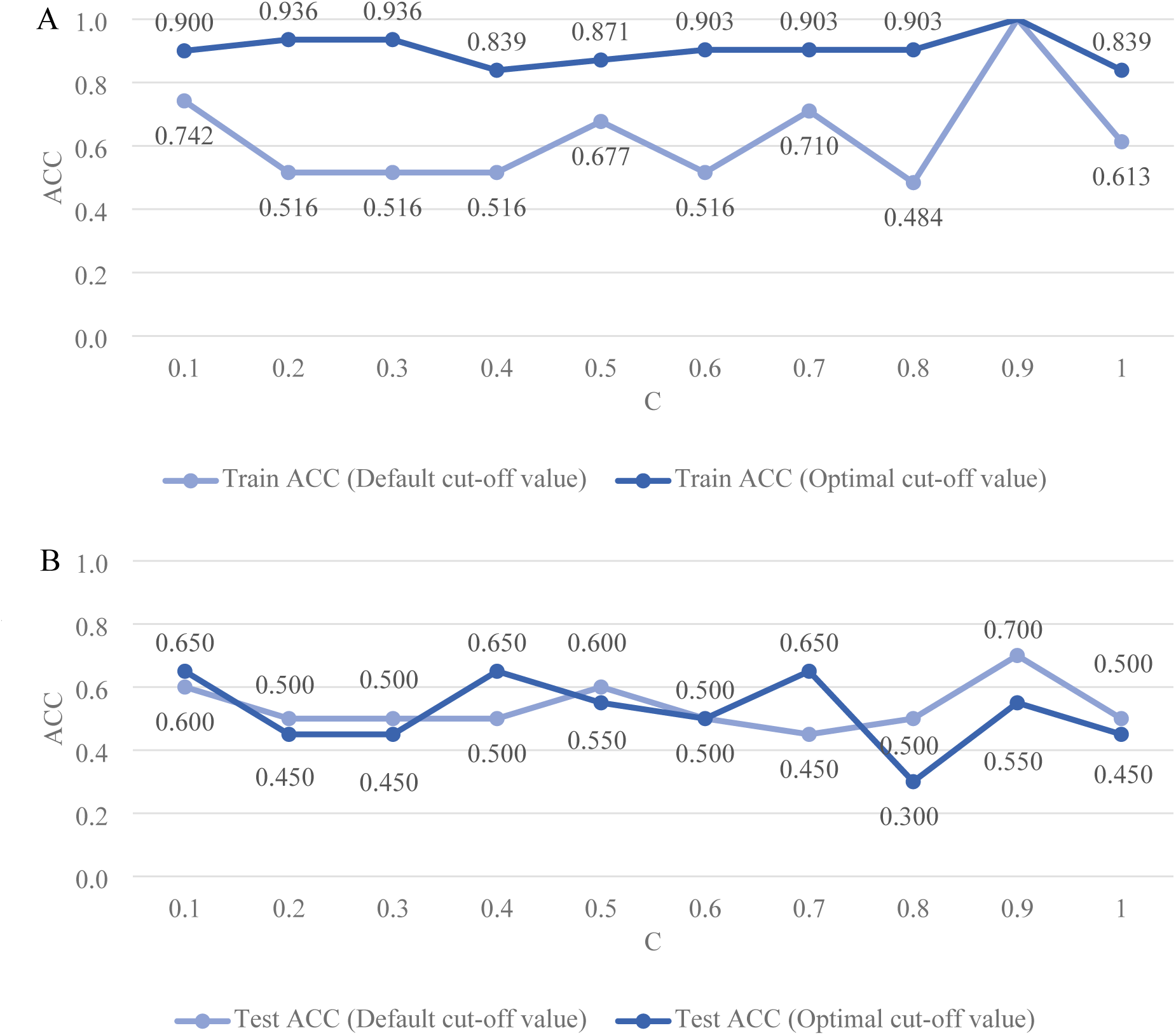
A. ACC for the training data set resulting from using the 0.5 cut-off (default) and the estimated cut-off (optimal) values. B. ACC for the test data set resulting from using the 0.5 cut-off (default) and the estimated cut-off (optimal) values.

Another classification metric important from the practical perspective is the sensitivity (Figure 5A) which reflects algorithm’s the ability to correctly classify an animal as mastitis susceptible. Another classification metric important from the practical perspective is the sensitivity (Figure 5A) which reflects algorithm’s the ability to correctly classify an animal as mastitis susceptible. Although a very high sensitivity, ranging from 0.750 (C=1.0) to 1.000 (C=0.1, C=0.3, and C=0.9) was reached for the training data sets, the sensitivity of test data classification was very low, except for one classification model for C=0.1 which obtained the high sensitivity of 0.800. On the other hand, the specificity (Figure 5B) of the test data classification (i.e. the ability to correctly classify an animal as mastitis-resistant) was generally higher than sensitivity. With that, it became evident that for most of the algorithms, the identification of resistant individuals is easier. The classification of the test data also reveals an interplay between both metrics, so models with high sensitivity, result in low specificity and the opposite.

**Fig 5.**
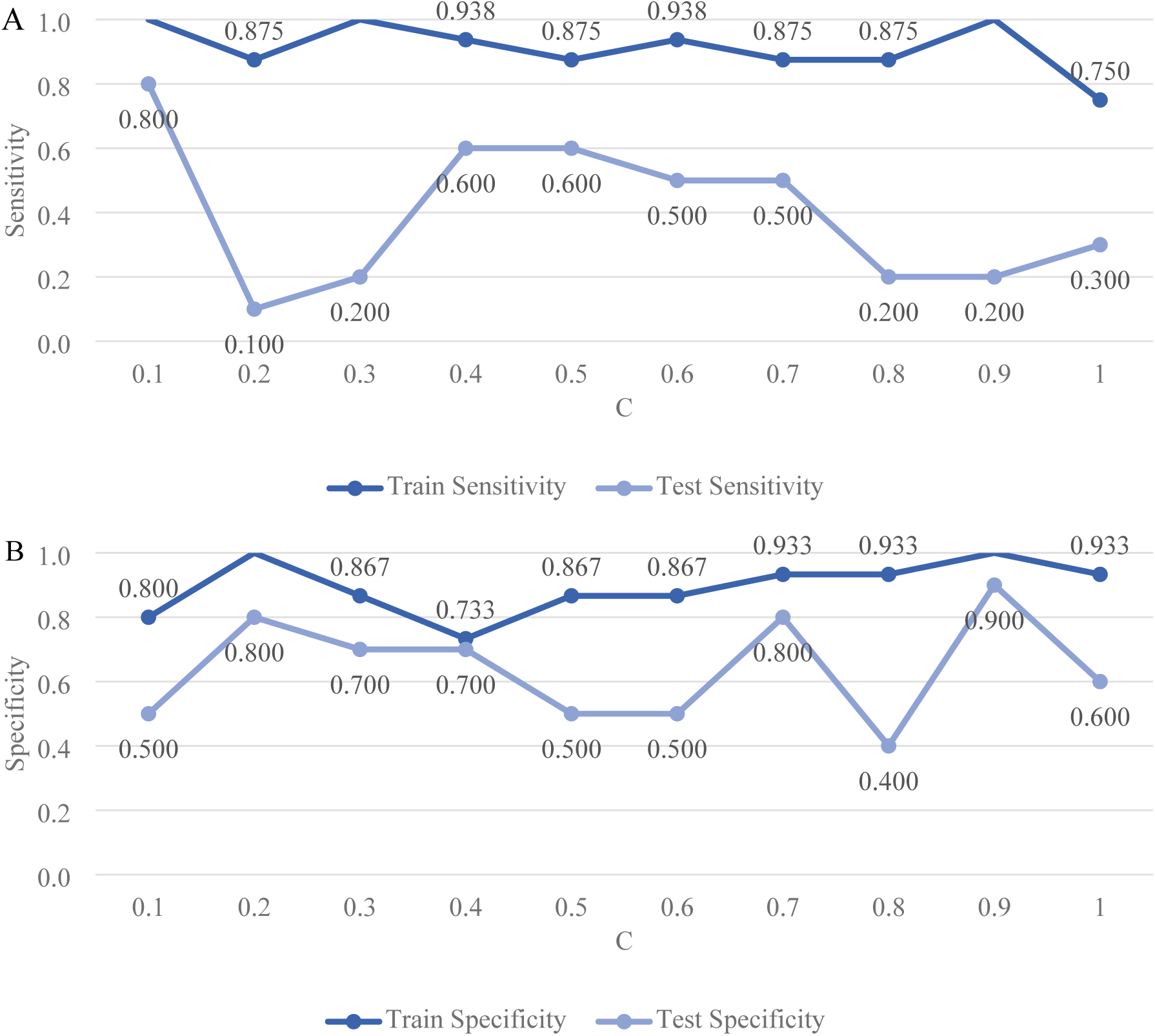
A. Classification sensitivity of train and test datasets based on the default cut-off values. B. Classification specificity of train and test datasets based on the default cut-off values.

By summarising the performance of the different DL architectures expressed by the AUC, accuracy, sensitivity, and specificity, the model corresponding to C=0.4, characterised by test AUC of 0.750, ACC equal to 0.650 (for the optimal cut-off), SENS=0.600, and SPEC=0.700 was selected as the best model and thus proceeded to genomic and functional annotations. This architecture utilises 204,642 SNPs.

### Genomic and functional annotation

From the 204,642 SNPs comprising the best model, 3,162 obtained significant (α = 0.05) SHAP values (Figure 6). 1,235 of those SNPs could be annotated to 966 genes – either by being located within their coding sequence (23 synonymous and 18 missense SNPs). Within non-coding sequences, the significant positions were located in introns (856 SNPs), non-coding transcripts (33 SNPs), non-coding exon variants (4 SNPs), upstream of genes (180 SNPs), downstream of genes (101 SNPs) 3’UTR variants (7 SNPs) or 5’UTR variants (13 SNPs). The remaining positions (1927SNPs) were intergenic variants. 746 of the SNPs were novel. Functions of the 966 genes were then tested for enrichment in GO terms from the biological process and the molecular function category, as well as in the KEGG and Reactome pathways. As a result, a single GO term related to the biological process (GO:0050804∼*modulation of chemical synaptic transmission* and thirteen GO terms related to the molecular function (GO:0005509∼*calcium ion binding*, GO:0030554∼*adenyl nucleotide binding*, GO:0032559∼*adenyl ribonucleotide binding*, GO:0005524∼*ATP binding*, GO:0032553∼*ribonucleotide binding*, GO:0032555∼*purine ribonucleotide binding*, GO:0017076∼*purine nucleotide binding*, GO:0005216∼*ion channel activity*, GO:0031267∼*small GTPase binding*, GO:0005096∼*GTPase activator activity*, GO:0017075∼*syntaxin-1 binding*, GO:0022838∼*substrate-specific channel activity*, GO:0008276∼*protein methyltransferase activity*) were significantly enriched, but none of GO terms describing cellular component, no KEGG and no Reactome pathway (S1 Table) The list presenting SNP assignment to the significant gene ontologies were given in the S2 Table.

**Fig 6.**
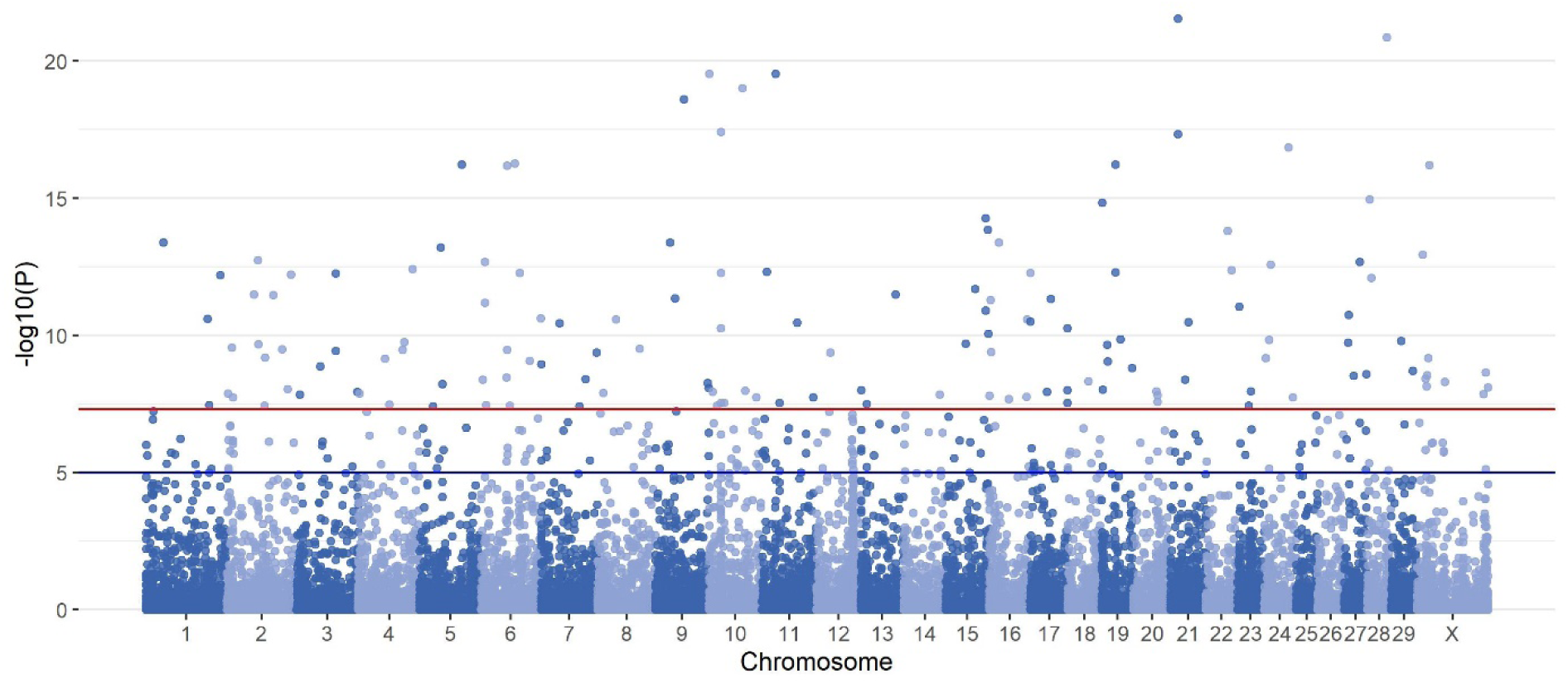
Manhattan plot with absolute values of 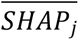 for the best classifying model. The red horizontal line represents the genome-wide significance threshold of p-value = 5.0 × 10^-8^ and the blue horizontal line represents the suggestive significance threshold of p-value = 1.0 × 10^-5^.

### GWAS for clinical mastitis in Polish Holstein-Friesian cows

In the association study of the large data set, 3,188 SNPs were significantly associated with clinical mastitis. The majority of those SNPs (99.72%) successively remapped to the ARS-UCD1.2 was annotated to 1209 genes, revealing 184 genes common with the genes marked by significant SNPs from the C=0.4 set. To further compare the functional annotation resulting from GWAS with that from the DL model, genes marked by significant SNPs from GWAS were tested for enrichment of GO terms. As a result, 11 GO terms from biological processes, 28 GO terms from molecular function, and 8 GO terms from cellular components were significantly enriched. Eight GO terms (GO:0005509∼*calcium ion binding*, GO:0030554∼*adenyl nucleotide binding*, GO:0032559∼*adenyl ribonucleotide binding*, GO:0005524∼*ATP binding*, GO:0032553∼*ribonucleotide binding*, GO:0032555∼*purine ribonucleotide binding*, GO:0017076∼*purine nucleotide binding*, GO:0005216∼*ion channel activity*, GO:0005096∼*GTPase activator activity*, GO:0022838∼*substrate-specific channel activity*) were overlapping between the GWAS-based significant enrichment and the DL-based functional enrichment (S1 Table).

## Discussion

Susceptibility to bovine mastitis is a complex trait since it is determined by a wide variety of bacteria, non-biological components (e.g. maintenance of milking equipment or post-milking teat disinfection) as well as by the genetic composition of an individual [20, 21]. Depending on the etiology the heritability of clinical mastitis varies between 0.01 to 0.25 [22], the latter indicating the considerable impact of the genetic component. In our study, using clinical mastitis as an example, we suggest and test a new three-step pipeline involving the bioinformatic, statistical and biological components to disentangle the functional component of a disease underlying a complex mode or inheritance.

### P>>n in classification

In a highly-dimensional data set, it is difficult to determine which of the explanatory variables are relevant [23] for prediction or classification. What is more, many of the explanatory variables are highly correlated so they do not provide unique information. Moreover, due to the number of features higher than the number of observations, it’s difficult not only to build the model because of overfitting but also to provide a concise and reproducible interpretation of its results, due to a galore of significant hits [24]. In the feature selection process, the originally very large number of explanatory variables (SNPs) was reduced to describe the response variable (mastitis-susceptible or mastitis-resistant), using the tuning parameter 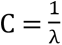 that regulates the sparsity of the estimator (i.e. the number of zero-valued coefficients), an approach similar to Fallerini et al. [25] that however seemed to use a single, predefined penalty value.

During this process, two major issues emerged. First, how to choose the number of SNPs (*N_SNP_*)? On the one hand, the lower *N_SNP_* makes the prediction model less expensive computationally, but on the other hand, it may result in a larger prediction error [26]. The problem can be extrapolated from genomics to database handling where selectivity estimation relates to estimating the number of records that satisfy query conditions [27]. However, the selectivity approach proposed in our study utilises quasi-empirical modelling of the LASSO penalty parameter λ by exploring the classification quality metrics of DL algorithms underlying a predefined range of penalty parameters that cover the full scope of potentially available SNPs. The second issue is the choice of the most appropriate measure of classification quality [28]. The ACC metric which is a standard in the evaluation of classification quality may become misleading when classification class sizes are imbalanced. Although in our data both class sizes were almost completely balanced, still the drawback exists that this classification quality metric relies on a binary assignment of individuals to TP/FP/TN/FN groups, regardless of the actual probability of classification that is the primary output of the sigmoid activation function from the last layer. To mitigate this problem, we proposed the exploration of the whole range of the probability parameter space (i.e. [0,1]) and the estimation of the cut-off value that guarantees the best ACC. Still, the approach based on the resampling of data requires a large sample size, which was not the case in our study. Another proposal is to evaluate the classification quality based on AUC that compares multiple thresholds of true positive (TPR) and false positive (FPR) rates [29] and provides a metric that simultaneously accounts for sensitivity and high specificity [30] of the classification.

Note that none of the compared DL architectures could be unequivocally classified as the best model by reaching the top scores of all the metrics applied. Practically, this means that there will be individuals which are incorrectly labelled as susceptible or resistant. So, the practical element of the best model selection is also to be driven by the interests of the end user of the classification, like milk producers in the case of our data or e.g. clinicians in the case of medical data.

### Functional interpretation of significant SNPs

All molecular functions significant in this study are fundamental in nearly every aspect of cell biology. According to Neculai-Valeanu and Ariton [31] mastitis causes alterations in the ionic dynamics of vascular components and it is mostly caused by the enormous cellular destruction and weakened milk–blood barrier. An increased ion concentration in milk during the mastitis infection involves sodium, potassium, calcium, magnesium, and chloride [32] which explains the overrepresented GOs related to ion binding and activity found in this study. The monoatomic ion channel activity (GO:0005216), according to the AmiGo definition [33], facilitates the diffusion of ions during their passage through a transmembrane aqueous pore or channel. On the other hand, the calcium ion binding ontology (GO:0005509) has been reported in the context of primary mammary epithelial cells (PMECs) infection, where the suppression of this ontological term was caused by Lipopolysaccharide (LPS), a toxin located in the outer membrane of Gram-negative bacteria [34]. Although this molecular function was suppressed, the authors observed that only a few ontological categories suppressed by LPS were significant and they hypothesized LPS induces immune, inflammatory and defence responses. In fact, immunological defence is a very energetically costly process requiring a shift in energy from less essential metabolic functions to the immune system in the presence of pathogens [35] which explains the role of ATP in the inflammation process. The ATP binding GO term (GO:0005524) was previously reported as the most representative molecular function in the context of infection with *Mycoplasma bovis*. This species is one of the major bovine pathogens that causes multiple diseases, including mastitis [36]. Moreover, because of the abovementioned ATP subtract and its specific channels, alterations in the amounts of molecules that activate these channels influence their activity (GO:0022838), causing in the case of bovine mastitis, an increase in the inflammatory response also indicated the role of the GTP binding in the immune response towards bovine mastitis which overlaps with our findings of the significant molecular functions of small GTPases binding (GO:0031267) and GTPase activator activity (GO:0005096). Moreover, GTPase-regulated pathways have often been mentioned in the mastitis context, especially concerning the inflammation caused by *Staphylococcus aureus*, but also *Mycoplasma bovis* [36–39]. Inflammation being the consequence of infection involves exocytosis which leads to the release of granule/vesicle contents to the cell exterior. It is of particular significance concerning tissue damage being the consequence of inflammatory cell activation and mediator elaboration [40]. Syntaxins are protein families playing an important role in exocytosis which may help to explain the syntaxin-1 binding molecular process (GO:0017075) found significant in this study. Regarding adenyl ribonucleotide binding (GO:0032559), not much has been reported in the mastitis-related literature. However, miRNA expression profiles were investigated in porcine mammary epithelial cells after contact with a potential mastitis-causing *Escherichia coli* strain. Predicted target genes of up- and downregulated miRNAs were significantly enriched in molecular functions including, among others, adenyl ribonucleotides [41]. Additionally, adenyl nucleotide binding (GO:0030554) and the ribonucleotide binding term (GO:0032553) were reported as significant on the altered molecular expression of the signalling pathway in mammary tissue of cattle with mastitis. The former was enriched in genes overexpressed in the mammary tissue of mastitis-infected cows [42]. Also, the nervous system plays a role in the response to infection since the immune system communicates with the nervous system to coordinate the immune response through signalling molecules, such as neurotransmitters and neuromodulators [43] which is reflected by the significant enrichment of the ontology of modulation of chemical synaptic transmission (GO:0050804). Purines are the bases of DNA and RNA that are required for the synthesis of nucleic acids, and thereafter proteins, and other metabolites, as well as for reactions that require energy [44]. The relation between nucleotide biosynthesis and bacterial pathogenesis in diseases was reported by Goncheva et al. [45] and demonstrates a connection with purine ribonucleotide and nucleotide binding ontologies (GO:0032555, GO:0017076) significant in our study. Finally, on the epigenetic level, Usman et al. [46] reported that the promoter regions of the JAK2 and the STAT5A genes were hypo-methylated in cows with mastitis which is in line with the significance of the protein methylation ontology (GO:0008276) estimated in our study. With this contribution, we emphasize the importance of exploiting multiple aspects of the bioinformatic analysis of biological data, which shall go beyond the application of bioinformatic software, but also comprise elements of feature preselection in the multidimensional data that are nowadays typical in genomics, multilevel model selection, statistical analysis and finally biological interpretation. This approach is especially important in the analysis of phenotypes with a complex mode of inheritance, like, in our case, clinical mastitis, that is influenced by multiple genes of varying effects and not necessarily by just a few genes of large effects. Moreover, due to low to moderate heritabilities, the effect of those genes is very likely also dependent on the environment (although this concept was not formally tested in our study).

## Materials and Methods

### Sequenced animals

52 Polish Holstein-Friesian cows were selected from the same herd with 991 clinical mastitis cases diagnosed by a veterinarian. All cows were kept in the same barn under unified conditions and were fed the same balanced diet. Moreover, they were born in the same year and season and also calved at the same age. The 52 individuals were divided into a *training group* of 32 cows and a *test group* of the remaining 20 cows. In the *training group,* cows were paternal half-sibs matched by the number of recorded parities, production level, and birth year but differed in their mastitis resistance status. So that 16 cows were mastitis-resistant and had no incidence of clinical mastitis throughout their production life, while 16 mastitis-susceptible cows underwent multiple disease incidences. Their genomic DNA was sequenced with the Illumina HiSeq2000 platform in the paired-end mode and with a read length of 100 bp. The number of raw reads generated for a single animal ranged between 164,984,147 and 472,265,620. The experimental design and the training dataset were described in detail by Szyda et al. [47]. The *test group* was composed of 10 mastitis-susceptible cows and 10 mastitis-resistant cows sequenced with the Illumina NovaSeq 6000 platform in the paired-end mode with 150 bp reads length. In this group, the number of reads available per individual ranged between 311,675,740 and 908,861,126.

### Genotyped animals

Another set of Polish Holstein-Friesian cows with clinical mastitis records was ascertained from the PLOWET database used for veterinarian-recorded health traits in four experimental dairy farms belonging to the National Institute of Animal Production. Among 1,499 individuals, clinical mastitis was recorded for 712 cows. Each cow was directly genotyped or imputed to the Illumina BovineSNP50K beadchip version 2. Genotype preprocessing comprised removing SNPs with a minor allele frequency below 0.01 and the quality of genotyping below 99% which resulted in 53,557 SNPs remaining for downstream analysis. Additionally, multigenerational pedigree records since 1914, comprising 8,944 ancestors of the genotyped cows were available for the estimation of their additive genetic relationship.

### Data processing scheme

The first part of the analysis, i.e., *the bioinformatic pipeline*, aims toward estimating the set of SNPs that forms the input for the DL-based classification scheme. Then, the *statistical pipeline* is used for the selection of the single best-classifying model comprising its underlying neural network architecture, hyperparameters, and the subset of SNPs and cut-off values estimations. Finally, the *biological pipeline* is imposed on significant SNPs from the best classifying model to provide the data set relevant biological explanation consisting of genome annotation and enrichment analysis. All protocols of the study are visualized in Figure 7.

**Fig 7.**
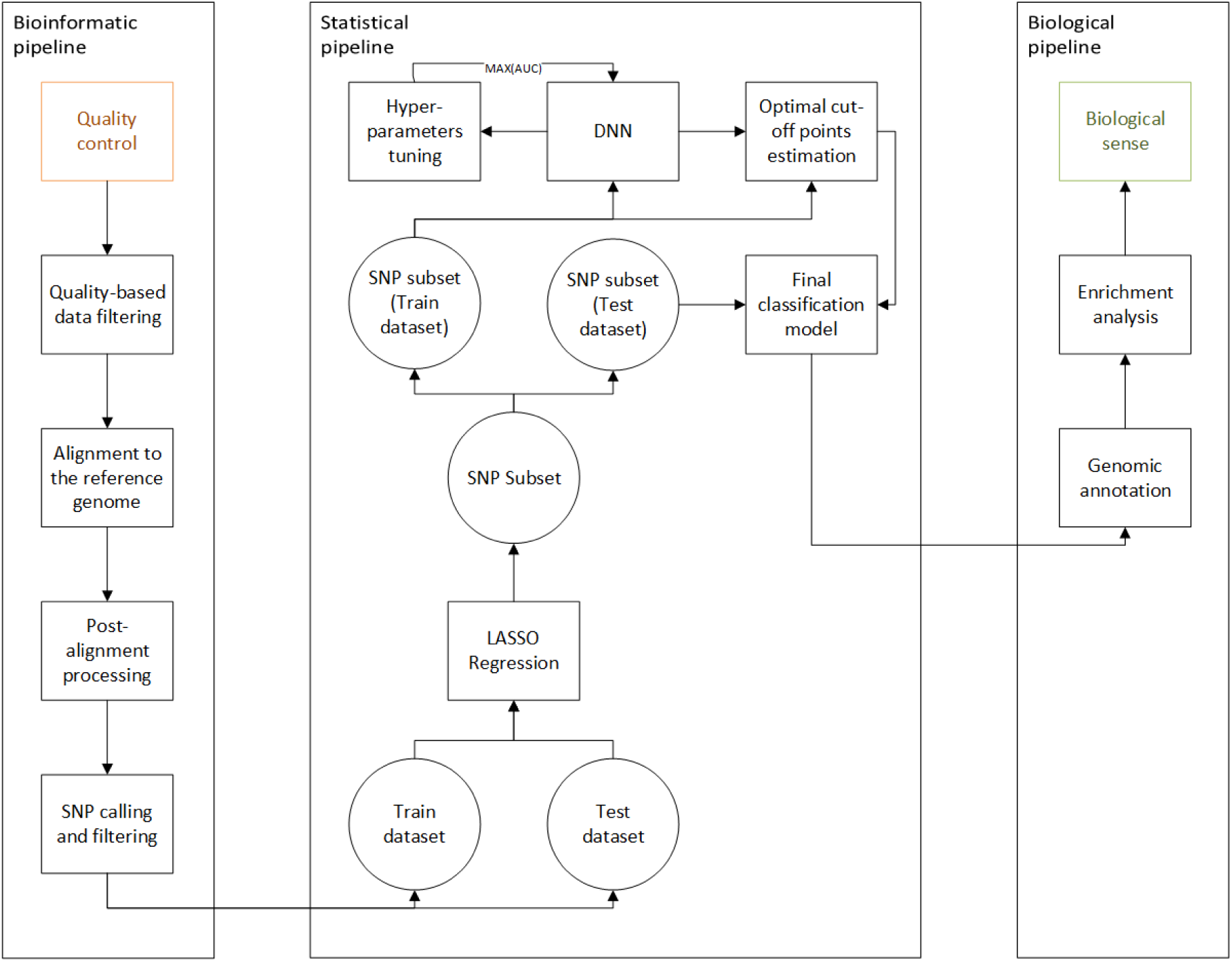
Flow diagram of data analysis.

### Bioinformatic pipeline

The bioinformatics pipeline for SNP identification consisted of: (i) the quality control step performed using FASTQC software [48], (ii) the quality-based raw data filtering step with Trimmomatic [49], (iii) the alignment to the ARS-UCD1.2 reference genome (NCBI accession number: PRJNA391427) with BWA-MEM [50], (iv) the post-alignment processing step with Samtools [51] and Bedtools [52] packages, (v) the SNP calling using GATK package [53], and (vi) the SNP filtering step with VCFtools [54]. In detail, based on the quality control report (i) the decision on trimming low-quality sequences was made. The procedure was performed by scanning each read with a 4-base sliding window and trimming it when the average of the 4-base qualities dropped below 20. The minimum length of read after trimming was set to 60 bp. The alignment of short reads to the reference genome was done with default parameters. Standard post-alignment processes included sorting and indexing of aligned sequences, PCR duplicates removal and further quality control. The variant calling step followed the best practice protocol provided by Van der Auwera and O’Connor [55]. Variant filtering incorporated removing variants with more than one alternative allele, identification quality below 20, and read depth at the variant site below 10. After all the above-mentioned edits, cows with average genome coverage below 7 were removed from downstream analyses.

### Statistical pipeline

#### the logistic LASSO regression

The first step, SNP preselection was performed by applying a logistic regression model:

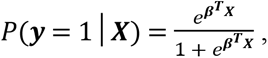

where, *P*(***y*** = 1|***X***) represents the probability of being mastitis-susceptible for each cow conditional on SNP genotypes ***X*,** with a LASSO [56] penalty (λ) imposed on the SNP effect estimator (***β̂***):

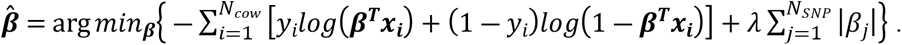

This logistic regression model was implemented with Python through the Scikit learn library [57] using the incremental gradient likelihood optimisation method with support for non-strongly convex composite objectives [58] and the L1 regularisation for SNP effect estimation. The penalty was expressed as 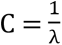, while various penalties were implemented using a grid over C within the interval (0.1; 1] with a step of 0.1. Note, that the smaller the λ, the more SNP estimates will be shrunk toward zero.

#### the deep learning algorithm

SNP sets pre-selected by LASSO with different penalties were further used in a deep learning classifier that was implemented via the Keras interface [59] with the TensorFlow [60] library in Python. The rectified linear unit (*ReLU*) function: *f*(*μ*) = max(0,*u*), was used as the activation function for all, except the last layer, for which the sigmoid function was applied 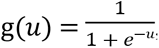, where *μ* is the node value calculated using the total summation of the input node values by their assigned. After each layer, the dropout regularization was applied resulting in a predefined fraction of the input units being set to zero. The Adam algorithm [61] implementing the stochastic gradient descent approach was used to optimise the binary cross-entropy loss function.

#### the hyperparameter tuning and validation

During the learning process, no fixed DL architecture was imposed, instead, the final architecture comprising the number of layers and neurons per layer, the dropout rate within each layer, and the learning rate for the optimisation algorithm was selected from the set of architectures dynamically sampled using the Optuna software [62] with one fixed hyperparameter *label smoothing* which transformed binary labels into probabilities. In particular, the *TPESampler* implemented in the Optuna was used for searching over DL algorithm hyperparameters with the number of iterations i.e. a single execution of hyperparameter estimation was set to 50. The pre-defined range of sampled hyperparameters was given in Table 2. which additionally summarises parameters that were set to fixed values, i.e. were not estimated, by the Optuna. A sampling of DL architectures was performed separately for each SNPs subset defined by different LASSO penalties (C). Additionally, the following mechanisms were implemented to prevent overfitting: (i) the early stopping mechanism, which terminates training of a given DL architecture when a value of the loss function did not decrease for 5 epochs, (ii) the pruning algorithm based on the Area Under the Curve (AUC) criterion, implemented via the *MedianPruner* method that terminates learning for DL architectures that result in small AUC. The training process was evaluated using a 4-cross-validation.

**Table 2.** Hyperparameters of the DL algorithm sampled by the Optuna software or treated as fixed.

| *Sampled hyperparameters* | *Range* |
| --- | --- |
| Number of layers | [1, 6] |
| Number of units per layer | [4, 50] |
| Dropout rate | [0.2, 0.4] |
| Learning rate | [1.0e-12, 1.0e-8] |
| *Fixed hyperparameters* |  |
| Number of epochs | 300 |
| Label smoothing | 0.2 |

#### the estimation of the optimal cut-off

The final DL architecture estimated for each LASSO penalty was applied to classify the test data set. For each individual, the output from the last layer, resulting from the sigmoid activation function, was expressed as a probability of being mastitis-susceptible. However, instead of applying a default 0.5 cut-off, for each of the selected DL architectures, the optimal probability cut-off was estimated using the cutpointR package [63] implemented in the R, based on the classification of cows from the training data set. In particular, the algorithm implemented into cutpointR determines the optimal cut-off value by maximizing the ACC metric. Estimates of the optimal cut-off values were obtained based on 1000 bootstrap samples of the training data set.

#### the selection of significant SNPs

For each SNP, SHAP values were used to assess the importance of each SNP on the classification:

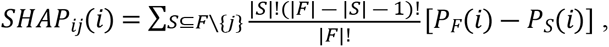

where *F* denotes a full set of SNPs, *S* is a subset of F with *j*-th SNP removed, *P_F_*(*i*) represents a probability of an *i*-th individual being mastitis-susceptible estimated based on full SNP set F and *P_S_*(*i*) represents a probability of an *i*-th individual being mastitis-susceptible estimated based on the subset S. Due to a very large number of SNPs, the SHAP values were not calculated directly, following the above formula, but were computed approximately using the DeepExplainer software [64]. The SHAP values were then rescaled to **z** scores: 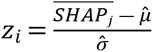, where 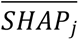 stands for the mean of *j*-th SNP across all individuals, 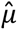 represents the mean and 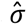 is a standard deviation of 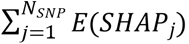 to assess each SNPs significance by testing (*H*0:*z_i_* ≤ 0 *vs*. *H*1:*z_i_* > 0) based on P-values from the standard normal distribution. Each p-value was transformed to a false discovery rate (FDR) [65] to account for multiple testing.

#### the evaluation of DL classifiers

The AUC metric was used as an indicator of the performance of each DL [66] while for the selection of the final, i.e. the best, classifier, sensitivity (SENS), specificity (SPEC) and accuracy (ACC) metrics were considered. Those metrics are based on the following classification outcomes:

i. True positive (TP), defined as the scenario when a mastitis-susceptible individual was classified as mastitis-susceptible,
ii. False positive (FP), defined as the scenario when a mastitis-resistant individual was classified as mastitis-susceptible,
iii. True negative (TN), defined as the scenario when a mastitis-resistant individual was classified as mastitis resistant,
iv. False negative (FN), defined as the scenario when a mastitis-susceptible individual was classified as mastitis-resistant,

and were defined as: 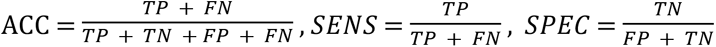. Among DL architectures with large AUC the best classifier was then chosen based on the highest accuracy, sensitivity, and specificity.

### Biological pipeline

SNPs from the best algorithm with significant (FDR<0.05) were annotated to genes from the ARS-UCD1.2 reference assembly by the Variant Effect Predictor (VEP) tool [67] considering the maximum upstream/downstream 5000 bp distance from the closest gene. Furthermore, for the genes marked by significant SNPs the enrichment analysis was performed by the Database for Annotation, Visualization and Integrated Discovery (DAVID) tool [68] using the most specific levels of Gene Ontologies [69] defined for Biological Processes (GO-BP), Molecular Functions (GO-MF), GO Cellular Component (CC) as well as for metabolic pathways defined by KEGG [70] and Reactome [71] databases.

### Genome-wide association study for clinical mastitis in genotyped cows

The association study was carried out using a multi-SNP approach based on the following mixed linear model:

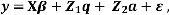

where ***y*** is a binary clinical mastitis status, ***β*** is a vector of fixed effect represented by a general mean and age at diagnosis, vector ***q*** contains SNP effects, ***a*** is a vector of additive polygenic effects of cows that were not explained by SNP genotypic variation, and vector ε contains error terms. ***Z***_1_ is a design matrix for SNP genotypes, which was parameterized as -1, 0, or 1 for a homozygous, heterozygous and an alternative homozygous SNP genotype, respectively. The covariance structure of the model was given by:

- 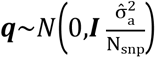, with I being an identity matrix, 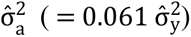 representing the additive genetic variance component and N_snp_ being equal to the number of SNPs (=53,557)
- 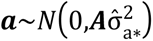, where ***A*** is the numerator relationship matrix calculated based on the pedigree relationship and 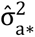 is the rest of additive genetic variance that was not explained by SNPs 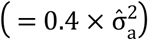,
- 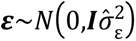 where ***I*** is an identity matrix and 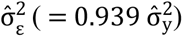 representing the residual variance.

The estimation of model parameters was based on solving the mixed model equations introduced by Henderson (1984):

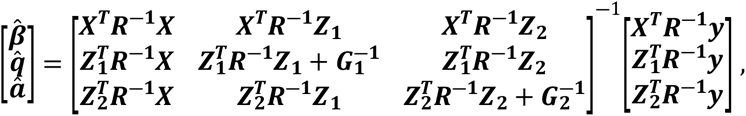

where 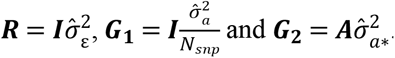. Consequently, the variance of y is then given by z_1_ 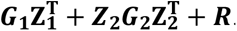.

For testing the hypotheses (*H*_0_:*q* = 0 vs. *H*_1_:*q* ≠ 0), we used the Wald test: 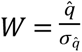, where 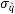 is a standard error of the estimated SNP effect *q*. Under H_0_ this statistic follows the standard normal distribution. The multiple testing correction was carried out via the Bonferroni approach (Dunnett, 1955). The positions were remapped from UMD3.1 to ARS-UCD1.2 reference genome using NCBI Genome Remapping Service [73] with default settings (minimum bases ratio for remapping=0.5 and maximum difference ratio between source and target length=2.0) and annotated to genes from the ARS-UCD1.2 reference assembly by the VEP tool.

## Acknowledgments

This work was supported by the project 2019/35/O/NZ9/00237 funded by The National Science Centre (NCN). Calculations were carried out at the Wroclaw Centre for Networking and Supercomputing and Poznan Supercomputing and Networking Center.

## Author contributions

KK: performed most of Data Curation, Formal Analysis including Validation, Visualisation as well as significantly contributed to Methodology and Writing of the Original Draft; MM: contributed to Data Curation and Formal Analysis as well as to Writing of the Original Draft, PB: contributed to the development of Methodology and to the Supervision of the research; KWM was responsible for the development and supervision of main Resources; TS: performed parts of the Formal Analysis and the underlying Methodology; PT: was responsible for the development and supervision of additional Resources; WJ: performed parts of the Formal Analysis; JS: was responsible for the Conceptualization, development of parts of the Methodology, as well as to Supervision and Funding Acquisition for the project.

## Supporting information

S1 Table. Significant GO terms with related gene names and SNP numbers inside them. The enrichment analysis was carried out on significant SNPs subset based on selected DL-based SNPs set. The GO terms overlapping between the GWAS-based and the DL-based significant enrichment have been underlined.

S2 Table. Significant SNPs from the best model with their associated SHAP values and annotation to gene ontologies with corresponding p-values.

## References

1. Routhier E, Mozziconacci J. Genomics enters the deep learning era. PeerJ. 2022;10: e13613. doi:10.7717/peerj.13613

2. Cao C, Liu F, Tan H, Song D, Shu W, Li W, et al. Deep Learning and Its Applications in Biomedicine. Genomics Proteomics Bioinformatics. 2018;16: 17–32. doi:10.1016/j.gpb.2017.07.003

3. European “1+ Million Genomes” Initiative. In: European Union [Internet]. Available: digital-strategy.ec.europa.eu/en/policies/1-million-genomes

4. Hayes BJ, Daetwyler HD. 1000 Bull Genomes Project to Map Simple and Complex Genetic Traits in Cattle: Applications and Outcomes. Annu Rev Anim Biosci. 2019;7: 89–102. doi:10.1146/annurev-animal-020518-115024

5. Cios KJ, Mamitsuka H, Nagashima T, Tadeusiewicz R. Computational intelligence in solving bioinformatics problems. Artif Intell Med. 2005;35: 1–8. doi:10.1016/j.artmed.2005.07.001

6. Asgari E, Mofrad MRK. Continuous Distributed Representation of Biological Sequences for Deep Proteomics and Genomics. Kobeissy FH, editor. PLoS One. 2015;10: e0141287. doi:10.1371/journal.pone.0141287

7. Liao JG, Chin K-V. Logistic regression for disease classification using microarray data: model selection in a large p and small n case. Bioinformatics. 2007;23: 1945–1951. doi:10.1093/bioinformatics/btm287

8. Severe Covid-19 GWAS Group, Ellinghaus D, Degenhardt F, Bujanda L, Buti M, Albillos A, et al. Genomewide Association Study of Severe Covid-19 with Respiratory Failure. N Engl J Med. 2020;383: 1522–1534. doi:10.1056/NEJMoa2020283

9. Zhao X, Qiao D, Yang C, Kasela S, Kim W, Ma Y, et al. Whole genome sequence analysis of pulmonary function and COPD in 19,996 multi-ethnic participants. Nat Commun. 2020;11: 5182. doi:10.1038/s41467-020-18334-7

10. Wesolowska-Andersen A, Zhuo Yu G, Nylander V, Abaitua F, Thurner M, Torres JM, et al. Deep learning models predict regulatory variants in pancreatic islets and refine type 2 diabetes association signals. Elife. 2020;9. doi:10.7554/eLife.51503

11. Sundaram L, Gao H, Padigepati SR, McRae JF, Li Y, Kosmicki JA, et al. Predicting the clinical impact of human mutation with deep neural networks. Nat Genet. 2018;50: 1161–1170. doi:10.1038/s41588-018-0167-z

12. Cheng L, Karkhanis P, Gokbag B, Liu Y, Li L. DGCyTOF: Deep learning with graphic cluster visualization to predict cell types of single cell mass cytometry data. Miller-Jensen K, editor. PLoS Comput Biol. 2022;18: e1008885. doi:10.1371/journal.pcbi.1008885

13. Bychkov D, Linder N, Turkki R, Nordling S, Kovanen PE, Verrill C, et al. Deep learning based tissue analysis predicts outcome in colorectal cancer. Sci Rep. 2018;8: 3395. doi:10.1038/s41598-018-21758-3

14. Lundberg SM, Lee S-I. A Unified Approach to Interpreting Model Predictions. Proceedings of the 31st International Conference on Neural Information Processing Systems. Red Hook, NY, USA: Curran Associates Inc.; 2017. pp. 4768–4777.

15. Ruegg PL. Investigation of mastitis problems on farms. Veterinary Clinics of North America: Food Animal Practice. 2003;19: 47–73. doi:10.1016/S0749-0720(02)00078-6

16. Halasa T, Huijps K, Østerås O, Hogeveen H. Economic effects of bovine mastitis and mastitis management: A review. Veterinary Quarterly. 2007;29: 18–31. doi:10.1080/01652176.2007.9695224

17. Jamali H, Barkema HW, Jacques M, Lavallée-Bourget E-M, Malouin F, Saini V, et al. Invited review: Incidence, risk factors, and effects of clinical mastitis recurrence in dairy cows. J Dairy Sci. 2018;101: 4729–4746. doi:10.3168/jds.2017-13730

18. Zhao X, Lacasse P. Mammary tissue damage during bovine mastitis: Causes and control1. J Anim Sci. 2008;86: 57–65. doi:10.2527/jas.2007-0302

19. Kossaibati MA, Esslemont RJ. The costs of production diseases in dairy herds in England. The Veterinary Journal. 1997;154: 41–51. doi:10.1016/S1090-0233(05)80007-3

20. Lakew BT, Fayera T, Ali YM. Risk factors for bovine mastitis with the isolation and identification of Streptococcus agalactiae from farms in and around Haramaya district, eastern Ethiopia. Trop Anim Health Prod. 2019;51: 1507–1513. doi:10.1007/s11250-019-01838-w

21. Smith KL, Hogan JS. Environmental Mastitis. Veterinary Clinics of North America: Food Animal Practice. 1993;9: 489–498. doi:10.1016/S0749-0720(15)30616-2

22. Nash DL, Rogers GW, Cooper JB, Hargrove GL, Keown JF, Hansen LB. Heritability of Clinical Mastitis Incidence and Relationships with Sire Transmitting Abilities for Somatic Cell Score, Udder Type Traits, Productive Life, and Protein Yield. J Dairy Sci. 2000;83: 2350–2360. doi:10.3168/jds.S0022-0302(00)75123-X

23. Asir D, Appavu S, Jebamalar E. Literature Review on Feature Selection Methods for High-Dimensional Data. Int J Comput Appl. 2016;136: 9–17. doi:10.5120/ijca2016908317

24. Simon R, Radmacher MD, Dobbin K, McShane LM. Pitfalls in the Use of DNA Microarray Data for Diagnostic and Prognostic Classification. JNCI Journal of the National Cancer Institute. 2003;95: 14–18. doi:10.1093/jnci/95.1.14

25. Fallerini C, Picchiotti N, Baldassarri M, Zguro K, Daga S, Fava F, et al. Common, low-frequency, rare, and ultra-rare coding variants contribute to COVID-19 severity. Hum Genet. 2022;141: 147–173. doi:10.1007/s00439-021-02397-7

26. Guyon I, Elisseeff A. An introduction to variable and feature selection. Journal of machine learning research. 2003;3: 1157–1182.

27. Hasan KMA, Siddique MdS, Rahman MdA. Selectivity estimation of large multidimensional data warehouses using logical grid directory. 2014 9th International Forum on Strategic Technology (IFOST). IEEE; 2014. pp. 9–13. doi:10.1109/IFOST.2014.6991060

28. Hicks SA, Strümke I, Thambawita V, Hammou M, Riegler MA, Halvorsen P, et al. On evaluation metrics for medical applications of artificial intelligence. Sci Rep. 2022;12: 5979. doi:10.1038/s41598-022-09954-8

29. Hand DJ. Measuring classifier performance: a coherent alternative to the area under the ROC curve. Mach Learn. 2009;77: 103–123. doi:10.1007/s10994-009-5119-5

30. Parikh R, Mathai A, Parikh S, Chandra Sekhar G, Thomas R. Understanding and using sensitivity, specificity and predictive values. Indian J Ophthalmol. 2008;56: 45. doi:10.4103/0301-4738.37595

31. Neculai-Valeanu A-S, Ariton A-M. Udder Health Monitoring for Prevention of Bovine Mastitis and Improvement of Milk Quality. Bioengineering. 2022;9: 608. doi:10.3390/bioengineering9110608

32. Kabelitz T, Aubry E, van Vorst K, Amon T, Fulde M. The Role of Streptococcus spp. in Bovine Mastitis. Microorganisms. 2021;9: 1497. doi:10.3390/microorganisms9071497

33. Carbon S, Ireland A, Mungall CJ, Shu S, Marshall B, Lewis S. AmiGO: online access to ontology and annotation data. Bioinformatics. 2009;25: 288–289. doi:10.1093/bioinformatics/btn615

34. Younis S, Javed Q, Blumenberg M. Meta-Analysis of Transcriptional Responses to Mastitis-Causing Escherichia coli. Hegde NR, editor. PLoS One. 2016;11: e0148562. doi:10.1371/journal.pone.0148562

35. Beiki H, Pakdel A, Javaremi AN, Masoudi-Nejad A, Reecy JM. Cattle infection response network and its functional modules. BMC Immunol. 2018;19: 2. doi:10.1186/s12865-017-0238-4

36. Chen S, Hao H, Zhao P, Ji W, Li M, Liu Y, et al. Differential Immunoreactivity to Bovine Convalescent Serum Between Mycoplasma bovis Biofilms and Planktonic Cells Revealed by Comparative Immunoproteomic Analysis. Front Microbiol. 2018;9. doi:10.3389/fmicb.2018.00379

37. Günther J, Petzl W, Bauer I, Ponsuksili S, Zerbe H, Schuberth H-J, et al. Differentiating Staphylococcus aureus from Escherichia coli mastitis: S. aureus triggers unbalanced immune-dampening and host cell invasion immediately after udder infection. Sci Rep. 2017;7: 4811. doi:10.1038/s41598-017-05107-4

38. Chen Y, Yang J, Huang Z, Yin B, Umar T, Yang C, et al. Vitexin Mitigates Staphylococcus aureus-Induced Mastitis via Regulation of ROS/ER Stress/NF-κB/MAPK Pathway. Gonçalves-de-Albuquerque CF, editor. Oxid Med Cell Longev. 2022;2022: 1–20. doi:10.1155/2022/7977433

39. Hughes K, Watson CJ. The Mammary Microenvironment in Mastitis in Humans, Dairy Ruminants, Rabbits and Rodents: A One Health Focus. J Mammary Gland Biol Neoplasia. 2018;23: 27–41. doi:10.1007/s10911-018-9395-1

40. Logan MR, Odemuyiwa SO, Moqbel R. Understanding exocytosis in immune and inflammatory cells: the molecular basis of mediator secretion. J Allergy Clin Immunol. 2003;111: 923–32; quiz 933. doi:12743551

41. Jaeger A, Hadlich F, Kemper N, Lübke-Becker A, Muráni E, Wimmers K, et al. MicroRNA expression profiling of porcine mammary epithelial cells after challenge with Escherichia coli in vitro. BMC Genomics. 2017;18: 660. doi:10.1186/s12864-017-4070-2

42. Wu J, Li L, Sun Y, Huang S, Tang J, Yu P, et al. Altered Molecular Expression of the TLR4/NF-κB Signaling Pathway in Mammary Tissue of Chinese Holstein Cattle with Mastitis. Srinivasula SM, editor. PLoS One. 2015;10: e0118458. doi:10.1371/journal.pone.0118458

43. Pavlov VA, Chavan SS, Tracey KJ. Molecular and Functional Neuroscience in Immunity. Annu Rev Immunol. 2018;36: 783–812. doi:10.1146/annurev-immunol-042617-053158

44. El Kouni MH. Purine Metabolism in Parasites: Potential Targets for Chemotherapy. Recent Advances in Nucleosides: Chemistry and Chemotherapy. Elsevier; 2002. pp. 377–416. doi:10.1016/B978-044450951-2/50013-8

45. Goncheva MI, Chin D, Heinrichs DE. Nucleotide biosynthesis: the base of bacterial pathogenesis. Trends Microbiol. 2022;30: 793–804. doi:10.1016/j.tim.2021.12.007

46. Usman T, Ali N, Wang Y, Yu Y. Association of Aberrant DNA Methylation Level in the CD4 and JAK-STAT-Pathway-Related Genes with Mastitis Indicator Traits in Chinese Holstein Dairy Cattle. Animals. 2021;12: 65. doi:10.3390/ani12010065

47. Szyda J, Frąszczak M, Mielczarek M, Giannico R, Minozzi G, Nicolazzi EL, et al. The assessment of inter-individual variation of whole-genome DNA sequence in 32 cows. Mammalian Genome. 2015;26: 658–665. doi:10.1007/s00335-015-9606-7

48. Andrews S. FastQC: A Quality Control Tool for High Throughput Sequence Data. 2010. Available: http://www.bioinformatics.babraham.ac.uk/projects/fastqc

49. Bolger AM, Lohse M, Usadel B. Trimmomatic: a flexible trimmer for Illumina sequence data. Bioinformatics. 2014;30: 2114–2120. doi:10.1093/bioinformatics/btu170

50. Li H, Durbin R. Fast and accurate short read alignment with Burrows–Wheeler transform. Bioinformatics. 2009;25: 1754–1760. doi:10.1093/bioinformatics/btp324

51. Li H, Handsaker B, Wysoker A, Fennell T, Ruan J, Homer N, et al. The Sequence Alignment/Map format and SAMtools. Bioinformatics. 2009;25: 2078–2079. doi:10.1093/bioinformatics/btp352

52. Quinlan AR, Hall IM. BEDTools: a flexible suite of utilities for comparing genomic features. Bioinformatics. 2010;26: 841–842. doi:10.1093/bioinformatics/btq033

53. McKenna A, Hanna M, Banks E, Sivachenko A, Cibulskis K, Kernytsky A, et al. The Genome Analysis Toolkit: A MapReduce framework for analyzing next-generation DNA sequencing data. Genome Res. 2010;20: 1297–1303. doi:10.1101/gr.107524.110

54. Danecek P, Auton A, Abecasis G, Albers CA, Banks E, DePristo MA, et al. The variant call format and VCFtools. Bioinformatics. 2011;27: 2156–2158. doi:10.1093/bioinformatics/btr330

55. der Auwera GA, O’Connor BD. Genomics in the cloud: using Docker, GATK, and WDL in Terra. O’Reilly Media; 2020.

56. Tibshirani R. Regression Shrinkage and Selection Via the Lasso. Journal of the Royal Statistical Society: Series B (Methodological). 1996;58: 267–288. doi:10.1111/j.2517-6161.1996.tb02080.x

57. Pedregosa F, Varoquaux G, Gramfort A, Michel V, Thirion B, Grisel O, et al. Scikit-Learn: Machine Learning in Python. J Mach Learn Res. 2011;12: 2825–2830.

58. Defazio A, Bach F, Lacoste-Julien S. SAGA: A Fast Incremental Gradient Method With Support for Non-Strongly Convex Composite Objectives. 2014. Available: http://arxiv.org/abs/1407.0202

59. Chollet F. Keras. 2015.

60. Abadi M, Barham P, Chen J, Chen Z, Davis A, Dean J, et al. Tensorflow: a system for large-scale machine learning. Osdi. 2016. pp. 265–283.

61. Kingma DP, Ba J. Adam: A Method for Stochastic Optimization. 2014. Available: http://arxiv.org/abs/1412.6980

62. Akiba T, Sano S, Yanase T, Ohta T, Koyama M. Optuna: A Next-generation Hyperparameter Optimization Framework. 2019. doi:1907.10902

63. Thiele C, Hirschfeld G. cutpointr : Improved Estimation and Validation of Optimal Cutpoints in R. J Stat Softw. 2021;98. doi:10.18637/jss.v098.i11

64. Shrikumar A, Greenside P, Kundaje A. Learning Important Features through Propagating Activation Differences. Proceedings of the 34th International Conference on Machine Learning - Volume 70. JMLR.org; 2017. pp. 3145–3153.

65. Benjamini Y, Hochberg Y. Controlling The False Discovery Rate - A Practical And Powerful Approach To Multiple Testing. J Royal Statist Soc, Series B. 1995;57: 289–300. doi:10.2307/2346101

66. Wu S, Flach P. A scored AUC Metric for Classifier Evaluation and Selection. 2005.

67. McLaren W, Gil L, Hunt SE, Riat HS, Ritchie GRS, Thormann A, et al. The Ensembl Variant Effect Predictor. Genome Biol. 2016;17: 122. doi:10.1186/s13059-016-0974-4

68. Sherman BT, Hao M, Qiu J, Jiao X, Baseler MW, Lane HC, et al. DAVID: a web server for functional enrichment analysis and functional annotation of gene lists (2021 update). Nucleic Acids Res. 2022;50: W216–W221. doi:10.1093/nar/gkac194

69. Ashburner M, Ball CA, Blake JA, Botstein D, Butler H, Cherry JM, et al. Gene Ontology: tool for the unification of biology. Nat Genet. 2000;25: 25–29. doi:10.1038/75556

70. Kanehisa M. KEGG: Kyoto Encyclopedia of Genes and Genomes. Nucleic Acids Res. 2000;28: 27–30. doi:10.1093/nar/28.1.27

71. Gillespie M, Jassal B, Stephan R, Milacic M, Rothfels K, Senff-Ribeiro A, et al. The reactome pathway knowledgebase 2022. Nucleic Acids Res. 2022;50: D687–D692. doi:10.1093/nar/gkab1028

72. Henderson CR. Applications of linear models in animal breeding. 1984.

73. Sayers EW, Bolton EE, Brister JR, Canese K, Chan J, Comeau DC, et al. Database resources of the national center for biotechnology information. Nucleic Acids Res. 2022;50: D20–D26. doi:10.1093/nar/gkab1112

